# Identifying Oscillatory Hyperconnectivity and Hypoconnectivity Networks in Major Depression Using Coupled Tensor Decomposition

**DOI:** 10.1101/2021.04.23.441123

**Authors:** Wenya. Liu, Xiulin. Wang, Jing. Xu, Yi. Chang, Timo. Hämäläinen, Fengyu. Cong

## Abstract

Previous researches demonstrate that major depression disorder (MDD) is associated with widespread network dysconnectivity, and the dynamics of functional connectivity networks are important to delineate the neural mechanisms of MDD. Cortical electroencephalography (EEG) oscillations act as coordinators to connect different brain regions, and various assemblies of oscillations can form different networks to support different cognitive tasks. Studies have demonstrated that the dysconnectivity of EEG oscillatory networks is related with MDD. In this study, we investigated the oscillatory hyperconnectivity and hypoconnectivity networks in MDD under a naturalistic and continuous stimuli condition of music listening. With the assumption that the healthy group and the MDD group share similar brain topology from the same stimuli and also retain individual brain topology for group differences, we applied the coupled nonnegative tensor decomposition algorithm on two adjacency tensors with the dimension of time × frequency × connectivity × subject, and imposed double-coupled constraints on spatial and spectral modes. The music-induced oscillatory networks were identified by a correlation analysis approach based on the permutation test between extracted temporal factors and musical features. We obtained three hyperconnectivity networks from the individual features of MDD and three hypoconnectivity networks from common features. The results demonstrated that the dysfunction of oscillation-modulated networks could affect the involvement in music perception for MDD patients. Those oscillatory dysconnectivity networks may provide promising references to reveal the pathoconnectomics of MDD and potential biomarkers for the diagnosis of MDD.

## I. Introduction

**M**AJOR depression disorder (MDD) is a globally common psychiatric disorder characterized by deficits of affective and cognitive functions [1]–[3]. It is almost a consensus to researchers that MDD is accompanied by abnormal functional connectivity (FC) between some brain regions, like cortical regions in the default mode network (DMN), rather than the aberrant response of individual brain regions [3]–[6]. An increasing amount of researches have demonstrated that FC presents the potential of temporal variability across different time-scales (from milliseconds to minutes) to support contineous cognitive tasks. This is termed as dynamic functional connectivity (dFC), and it represents the processes by which networks and subnetworks coalesce and dissolve over time, or cross-talk between networks [7]–[9]. Recently, researches have reported abnormal dFC of specific brain regions and neural networks in MDD using resting-state functional Magnetic Resonance Imaging (RS-fMRI) [3], [5], [9], [10]. For example, Demirtas *et al*. found a decreased variability of FC in the connections between the DMN and the frontoparietal network [5]. Kaiser *et al*. showed that MDD patients presented decreased dFC between medial prefrontal cortical (MPFC) regions and regions of parahippocampal gyrus within the DMN, but increased dFC between MPFC regions and regions of insula. They showed that MDD was related to abnormal patterns of fluctuating communication among brain systems involved in regulating attention and self-referential thinking [9]. The decreased dFC variability was reported between anterior DMN and right central executive network (CEN) in MDD, which indicated a decreased information processing and communication ability [10]. Existing researches about dFC in MDD mostly focus on resting-state conditions. However, little is known about the abnormalities of dFC during music listening conditions, especially when music therapy has become an attractive tool for MDD treatment [11]–[13].

Benefiting from the high temporal resolution, EEG can record electrical brain activity dynamics at a millisecond scale with rich frequency contents. The EEG oscillation acts as a bridge to connect different brain regions with resonant communication, which can regulate changes of neuronal networks and cause qualitative transitions between modes of information processing [14]–[16]. Impaired coordination of brain activity associated with abnormal electrophysiological oscillations contributes to the generation of psychaitric disorders [17]. Numerous studies have investigated EEG oscillatory FC of MDD in resting-state, and dysconnectivity networks are mostly notable in theta, alpha and beta oscillations [15], [18], [19]. However, most previous studies filter EEG signals into a range of frequency bands (e.g., 8-13 Hz for the alpha band), which considers the FC as a static state in a predefined frequency range and ignores exhaustive spectral dynamics in FC. Music perception is a complex cognitive task, which is characterized by dynamics of frequency-specific brain networks for musical features processing [20]–[24]. To the best of our knowledge, the oscillatory dFC in MDD during music perception has not been investigated yet.

Considering the temporal and spectral dynamics of spatial couplings (e.g., functional connectivity) for multiple participants in a cognitive task, a multi-way dataset structure is naturally formed. This multi-dimensional nature points to the adoption of tensor decomposition models instead of matrix decomposition models which normally fold some dimensions and ignore the hidden interactions across different modes [23], [25]–[28]. Canonical Polyadic (CP) decomposition is derived in terms of the sum of multiple rank-one tensors, and each rank-one tensor represents the covariation of the corresponding components from each mode [29], [30]. The CP decomposition is well implemented into the extraction of multi-mode EEG features from the multiway dataset (e.g., channel × frequency × time × subject) [30]–[33]. Recently, Zhu *et al*. applied CP decomposition to explore the task-related dFC characterized by spatio-temporal-spectral modes of covariation from the adjacency tensor (connectivity × time-subject × frequency) [22], [34]. However, those applications only focus on the decomposition of one single tensor, which are based on the assumption that the underlying spatio-spectral features are consistent among subjects or groups [24], [28]. Coupled tensor decomposition (CTD), the extension of tensor decomposition to multiple block tensors, enables the simultaneous extraction of common features shared among tensors and individual features specified for each tensor. For biomedical data, the coupled matrix, matrix-tensor or tensor decomposition (also known as linked component analysis) are mostly used for data fusion [35]–[37]. However, to the best of our knowledge, no studies have used CTD to investigate the pathologic networks of MDD or other psychiatric disorders.

In our study, we applied a low-rank double-coupled nonnegative tensor decomposition (DC-NTD) model to explore the temporal and spectral dynamics of spatial couplings in MDD during music listening. We collected the ongoing EEG data from healthy participants and MDD participants under music listening conditions, and projected the data from sensor space to cortical source space. Phase couplings were calculated by the phase lag index (PLI) along both temporal and spectral dynamics through time-frequency representations. Then, we constructed two fourth-order tensors of time × frequency × connectivity × subject for the healthy group and the MDD group. Considering the high computation load, the nonnegativity of the tensors (constrained to [0,1] due to PLI index) and high correlations in spatial and spectral modes, we applied a low-rank DC-NTD model which was more flexible to add desired constraints. Correlation analysis was conducted between temporal courses of brain activities and temporal courses of musical features to obtain the music-induced dFC. Finally, we summarized three oscillatory hypoconnectivity networks from common music-induced features which were active in the healthy group but not in the MDD group, and we also summarized three oscillatory hyperconnectivity networks from music-induced features specified in MDD. Figure 1 shows the diagram of the analysis pipeline of this study.

**Fig. 1:**
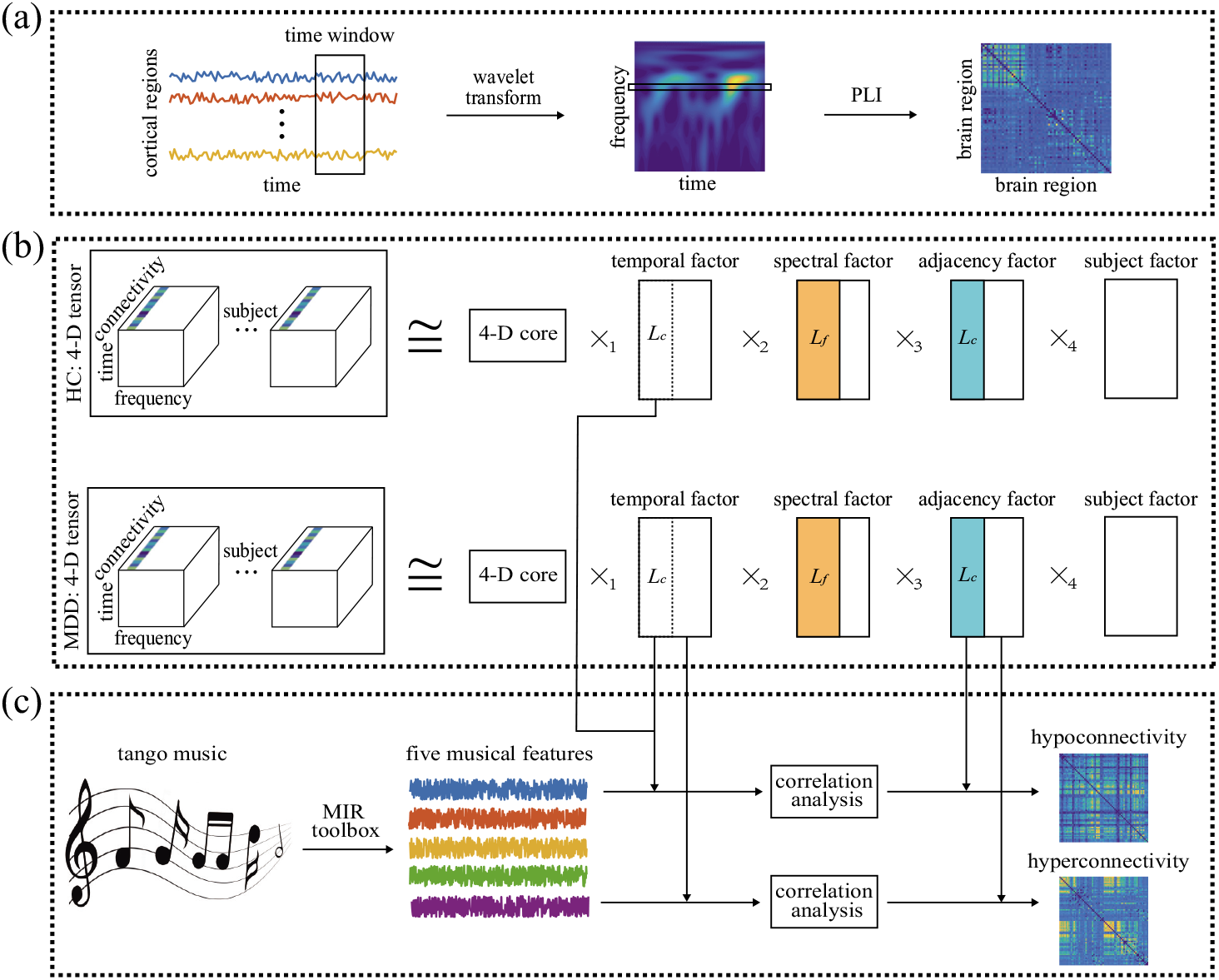
Diagram of the analysis pipeline. (a) Adjcency matrix construction in each time window and each frequency bin. After source reconstruction, the cortical signals were segmented by overlapping time windows, and wavelet transform was applied for each time course within each time window. Phase lag index was used to obtain the adjacency matrix for each time window and each frequency bin. (b) Adjacency tensor construction and decomposition. A 4-D adjacency tensor was constructed for each group with the dimension of time × frequency × connectivity × subject, and coupled tensor decomposition was implemented with coupled constraints in spectral and adjacency modes. The 4-D core tensor is superdiagonal with values of 1. (c) The identification of hyperconnectivity and hypoconnectivity networks by music modulation. Five musical features were extracted with MIR toolbox from tango music, and correlation analysis was conducted between musical features and decomposed temporal factors to identify music-induced brain networks. Hyperconnectivity and hypoconnectivity networks were summarized from the results of music modulation.

In this paper, scalars, vectors, matrices and tensors are denoted by lowercase, boldface lowercase, boldface uppercase and boldface script letters, respectively, e.g., *x*, ***x, X, 𝒳***. Indices range from 1 to their capital version, e.g., *i* = 1, …, *I*.

## II. Materials and methods

### A. Data description

#### 1) Participants

Twenty MDD patients and nineteen healthy controls (HC) participated in this experiment. All the patients were from the First Affiliated Hospital of Dalian Medical University in China. This study has been approved by the ethics committee of the hospital, and all participants signed the informed consent before their enrollment. None of the participants has reported hearing loss and formal training in music. All the MDD patients were primarily diagnosed by a clinical expert and tested according to Hamilton Rating Scale for Depression (HRSD), Hamilton Anxiety Rating Scale (HAMA) and Mini-Mental State Examination (MMSE). The means and standard deviations (SD) of age, gender, education and clinical measures for both groups were listed in Table I.

**TABLE I:** Means and standard deviations of age, gender, education and clinical measures for HC group and MDD group

|  | HC group |  | MDD group |  | p-value |
| --- | --- | --- | --- | --- | --- |
|  | Mean | SD | Mean | SD |  |
| Age | 38.4 | 11.8 (24-65) | 42.9 | 11.0(23-58) | >0.05 |
| Education | 13.6 | 3.8(6-20) | 12.8 | 3.4(6-16) | >0.05 |
| HRSD | 2.4 | 1.3(0-4) | 23.3 | 3.6(16-28) | <0.01 |
| HAMA | 2.4 | 1.3(0-5) | 19.2 | 3.0(15-25) | <0.01 |
| MMSE | 28.2 | 0.9(27-30) | 28.1 | 1.1(26-30) | >0.05 |
| Duration | 0 | 0 | 12.8 | 8.5(2-36) | - |
| Gender | 14 females, 5 males |  | 14 females, 6 males |  | - |

Abbreviations: HC, healthy controls; MDD, major depression disorder; SD, standard deviations; HRSD, Hamilton Rating Scale for Depression; HAMA, Hamilton Anxiety Rating Scale; MMSE, Mini- Mental State Examination.

#### 2) EEG data

During the experiment, participants were told to sit comfortably in the chair and listen to a piece of music. A 512-second musical piece of modern tango *Adios Nonino* by Astor Piazzolla was used as the stimulus due to its rich musical structure and high range of variation in musical features such as dynamics, timbre, tonality and rhythm [20], [38]. The EEG data were recorded by the Neuroscan Quik-cap device with 64 electrodes arranged according to the international 10-20 system. The electrodes placed at the left and right earlobes were used as the references.

The data were visually checked to remove obvious artifacts from head movement and down-sampled to *fs* = 256 Hz for further processing. Then 50 Hz notch filter and high-pass and low-pass filters with 1 Hz and 30 Hz cutoff were applied. Eye movements artifacts were rejected by independent component analysis (ICA).

#### 3) Musical features

In this study, two tonal and three rhythmic features were extracted by a frame-by-frame analysis approach using MIR toolbox [39]. The duration of each frame was 3 seconds, and the overlap between two adjacency frames was 2 seconds. Therefore, we got 510 samples for the time courses of each musical feature at a sampling frequency of 1 Hz. In this study, we only used the first *T* = 500 samples of each musical feature due to the length of recorded EEG data. Tonal features include Mode and Key Clarity, which represent the strength of major of minor mode and the measure of tonal clarity, respectively. Rhythmic features include Fluctuation Centroid, Fluctuation Entropy, and Pulse Clarity. Fluctuation centroid is the geometric mean of the fluctuation spectrum representing the global repartition of rhythm periodicities within the range of 0–10 Hz. Fluctuation Entropy is the Shannon entropy of the fluctuation spectrum representing the global repartition of rhythm periodicities. Pulse Clarity is an estimate of clarity of the pulse.

### B. Source reconstruction

Source reconstruction procedure was performed with open-source Brainstorm software [40]. For forward modeling, we used the symmetric boundary element method (BEM) to compute the volume-conductor model with the MNI-ICBM152 template corresponding to a grid of 15000 cortical sources. For source modeling, minimum norm estimate (MNE) was applied with a measure of the current density map and constrained dipole orientations (normal to cortex). Then, the Desikan-Killiany anatomical atlas was used to parcellate the cortical surface into *C* = 68 regions, and the principal component analysis (PCA) method was performed to construct the time course for each brain region.

### C. Dynamic functional connectivity

Many studies have reported that the communication of brain regions or neural populations depends on phase interactions for electrophysiological neuroimaging techniques, like EEG [41]. To avoid source leakage, the pairwise synchronization was estimated by PLI to map the whole-brain FC [42]. In this study, to assess the dFC across both time and frequency, we segmented the source-space data into *W* = 500 windows by the sliding window technique with a window length of 3 s and an overlap of 2 s according to the extraction framework of musical features. Then, we computed the time-frequency (TF) decomposition within each time window by the continuous wavelet transform with Morlet wavelets as basis function. We set the frequency bins as 0.5 Hz, and obtained *F* = 59 samples in frequency domain in the range of 1-30 Hz.

For the time window *w*, we can get the complex TF representation 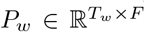 from wavelet transform, where *T*_*w*_ = 3*fs*, and the time and frequency-dependent phase at time *t*_*w*_ and frequency *f* can be obtained by

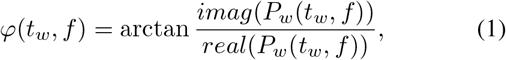

where *imag*() and *real*() represent the imaginary part and the real part of a complex value, respectively. For brain regions *I* and *j*, PLI can be computed as

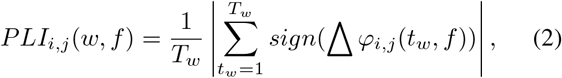

where, Δ *φ* _*i,j*_ (*t*_*w*_,*f*))= *φ* _*i*_ (*t*_*w*_,*f*) − *φ* _*j*_ (*t*_*w*_,*f*) is the phase difference of brain regions *i* and *j* at time *t*_*w*_ and frequency *f* in time window *w*. Therefore, for each time window and each frequency point, we can form an adjacency matrix ***A*** ∈ ℝ ^*C*×*C*^, where *C* means the number of brain regions. Because of the symmetry of FC matrix, we took the upper triangle of ***A*** and vectorized it to ***a*** ∈ ℝ ^*N* ×1^, where *N* = *C*(*C* − 1)/2 = 2278 represents the number of unique connections. Then, we can construct two adjacency tensors with the dimension of time × frequency × connectivity × subject, 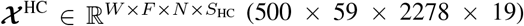 for the HC group and 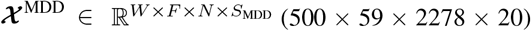 for the MDD group, where *S*_HC_ = 19 and *S*_MDD_ = 20 mean the number of subjects in the HC group and the MDD group, respectively.

### D. The application of low-rank coupled tensor decomposition

#### 1) Low-rank coupled tensor decomposition

With the constructed tensors 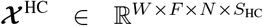 and 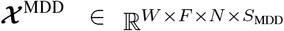 the corresponding CP decomposition can be represented as 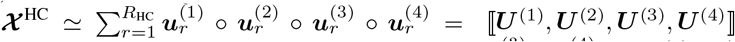 and 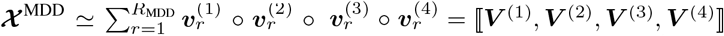 where ∘ denotes the vector outer product. 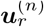 and 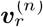 denote the *r*th component of factor matrices ***U*** ^(*n*)^ and ***V*** ^(*n*)^, *n* = 1, 2, 3, 4, in the modes of time, frequency, connectivity and subject for two groups. *R*_HC_ and *R*_MDD_ are the ranks of *𝒳*^HC^ and *𝒳*^MDD^, respectively. Considering the nonnegativity of constructed tensors and the coupled constraints in spectral and adjacency modes, we formulate it as a double-coupled nonnegative tensor decomposition (DC-NTD) model, where *𝒳*^HC^ and *𝒳*^MDD^ can be jointly analyzed by minimizing the following objective function:

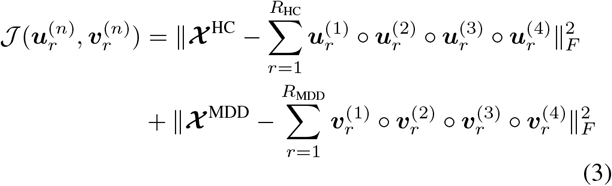

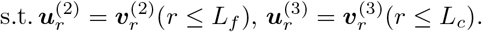

‖ · ‖_*F*_ denotes the Frobenius norm. *L*_*f*_ and *L*_*c*_ denote the number of components coupled in spectral and adjacency modes, and *L*_*f,c*_ ≤ min(*R*_HC_, *R*_MDD_). The fast hierarchical alternative least squares (FHALS), an accelerated version of the hierarchical alternative least squares (HALS) algorithm, has been effectively applied to a number of (coupled) tensor decomposition problems [24], [43], [44]. In this study, we apply the FHALS algorithm to optimize the DC-NTD problem in (3), and introduce the low-rank approximation to reduce computational complexity [45], [46].

Through the FHALS algorithm, the minimization problem in (3) can be converted into max(*R*_HC_, *R*_MDD_) rank-1 subproblems, which can be solved sequentially and iteratively. We take the *r*th subproblem as an example:

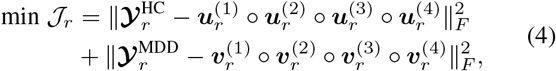

where 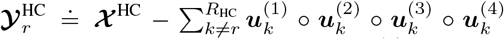 and 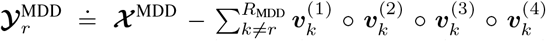.When calculating one of the variables, we need to fix the other variables and let the corresponding derivative be zero. For example, to determine 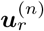, we let 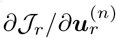 be zero, and then we can obtain the following solution:

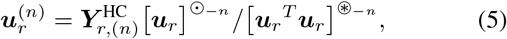

where 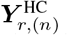 is the mode-*n* matricization of 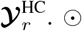 and ⊛ denote the Khatri-Rao product and Hadamard (element-wise) product. 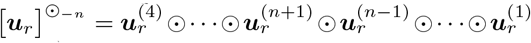 and 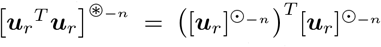 Taking 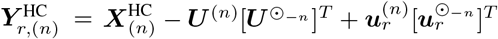 into (5), we can get

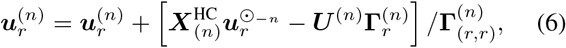

where 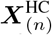 is the mode-*n* matricization of ***𝒳*** ^HC^ and **Г**^(*n*)^ =[***U*** ^*T*^ ***U***] ^⊛ −*n*)^ .Suppose that the rank-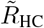 approximation of ***𝒳*** ^HC^ obtained by unconstrained tensor factorization is expressed as ⟦ *Ũ* ^(1)^, *Ũ* ^(2)^, *Ũ* ^(3)^, *Ũ* ^(4)^ ⟧,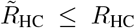 thus the mode-*n* unfolding of ***𝒳*** ^HC^ can be written as 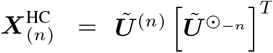 Therefore, the learning rule of 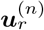 can be reformulated as follows:

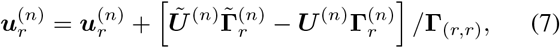

Where 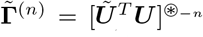. Analogously, we can obtain the learning rule of 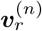 as follows:

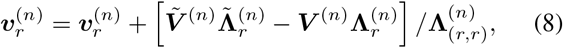

where **Λ**^(*n*)^ = [***V*** ^*T*^ ***V*]** ^⊛ −*n*)^ and 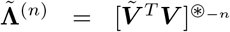 The rank-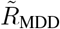 approximation of ***𝒳*** ^MDD^ is expressed as 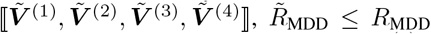 Specially, if 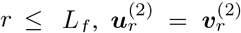 and if 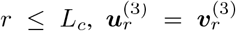, thus their solutions should be calculated as:

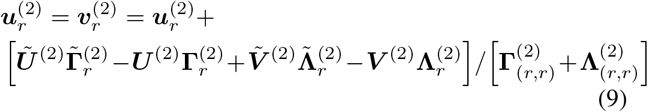

and

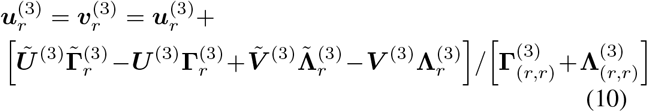

In order to obtain the nonnegative components, a simple “half-rectifying” nonlinear projection is applied. We update 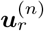 and 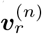 successively in each subproblem, and the max(*R*_HC_, *R*_MDD_) subproblems are optimized alternatively one after another until convergence. In this study, we adopt alternating least squares (ALS, [47]) algorithm to perform low-rank approximation. The FHALS-based DC-NTD algorithm is summarized in **Algorithm** 1.

##### Algorithm 1 DC-NTD-FHALS algorithm

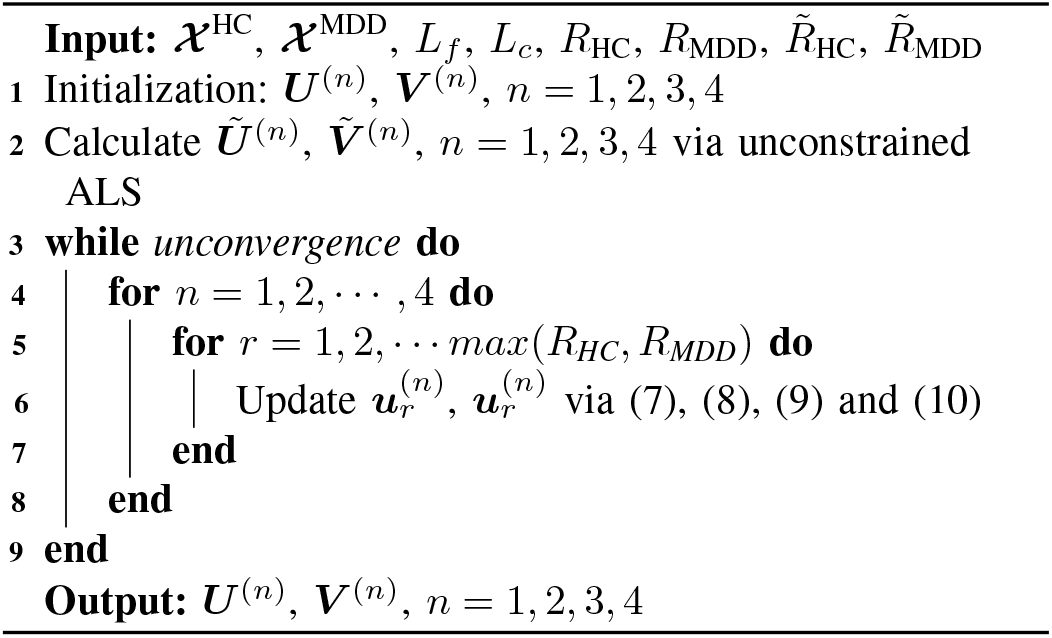

### 2) Selection of components

In this section, we will describe how to determine the number of totally extracted components *R*_*HC*_ and *R*_*MDD*_, which refers to the hidden information in low-dimensional space for each block data, and the number of coupled components *L* _*f*_ and *L* _*c*_, which reveal the common features between two-block data. For the selection of *R*_*HC*_ and *R* _*MDD*_, we performed PCA on the matricization data *𝒳* _(3)_ ∈ ℝ ^*F*×*WNS*^ unfolded along frequency mode for each block data, and kept the number of components with 95% explained variance. The selection of the number of coupled components is a key issue for the conduction of the DC-NTD-FHALS algorithm and the explanation of results, and it always becomes an open issue depending on practical applications. In this study, we performed the fourth-order CP tensor decomposition based on the FHALS algorithm on two-block data separately, and calculated the correlation maps of extracted components between two-block data in spectral and adjacency modes, respectively. According to the correlation maps, we will select the number of highly correlated (coupled) components. The detailed implication procedure will be described in the results section.

### E. Identification of music-induced hyperconnectivity and

After the conduction of the low-rank DC-NTD-FHALS algorithm, we need to identify the music-induced oscillatory networks that are abnormal involved in the MDD group. We conducted a correlation analysis approach between temporal factors and five musical features with Pearson correlation based on the permutation test method. To ensure the statistical significance of the correlation and consider the problem of multiple comparison, the Monte Carlo method and permutation test were applied to compute a significant threshold of correlation for each musical feature [20]–[22], [24]. For the time course of each musical feature, we kept the real part and replaced the imaginary part with random uniformly distributed phases, and performed Pearson correlation with the time courses of the extracted temporal components. Then, we repeated this procedure 100000 times, and obtained the threshold for each musical feature at a significant level of *p*_*corrected*_ < 0.05.

The coupled spectral and adjacency components are common features between CON and MDD groups, and the remaining components are individual features of each group. The oscillatory networks among common features that are involved in music perception in the HC group but not in the MDD group are identified as hypoconnectivity networks, and the oscillatory networks among individual features that are involved in music perception in the MDD group are identified as hyperconnectivity.

## III. Results

### A. Simulation validation

#### 1) simulation data

To validate the feasibility of the proposed method, we firstly applied it on the simulated data. Two tensors with the size of 500 × 59 × 2278, representing time × frequency × connectivity, were created as follows:

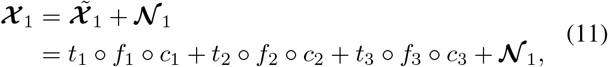

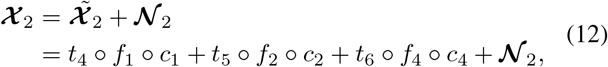

where 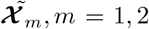 represented the ground truth networks, and **𝒩** _*n*_, *n* = 1, 2 were the nonnegative noise created by the absolute values of white noise with the size of 500 × 59 × 2278. In the time domain, each temporal component *t*_*i*_, *i* = 1, 2, …,6 was simulated by the absolute value of white noise to ensure the nonnegativity of the synthetic tensor ***𝒳***, and no coupled temporal component existed between two tensors. In the frequency domain, four spectral components *f*_*j*_, *j* = 1, 2, …, 4 were constructed by Hanning windows and white noise with bandwidth centered at 5 Hz, 10 Hz, 15 Hz and 20 Hz, and two spectral components were set to be coupled between two tensors. In the adjacency domain, four adjacency components *c*_*k*_, *k* = 1, 2, …, 4, representing auditory network (AUD), visual network (VIS), salience network (SAN), and dorsal attentional network (DAN), were constructed with the Desikan-Killiany anatomical atlas according to Kabbara’s work [48], and two adjacency components were coupled between two tensors. The synthetic data were shown in Figure 2(a).

**Fig. 2:**
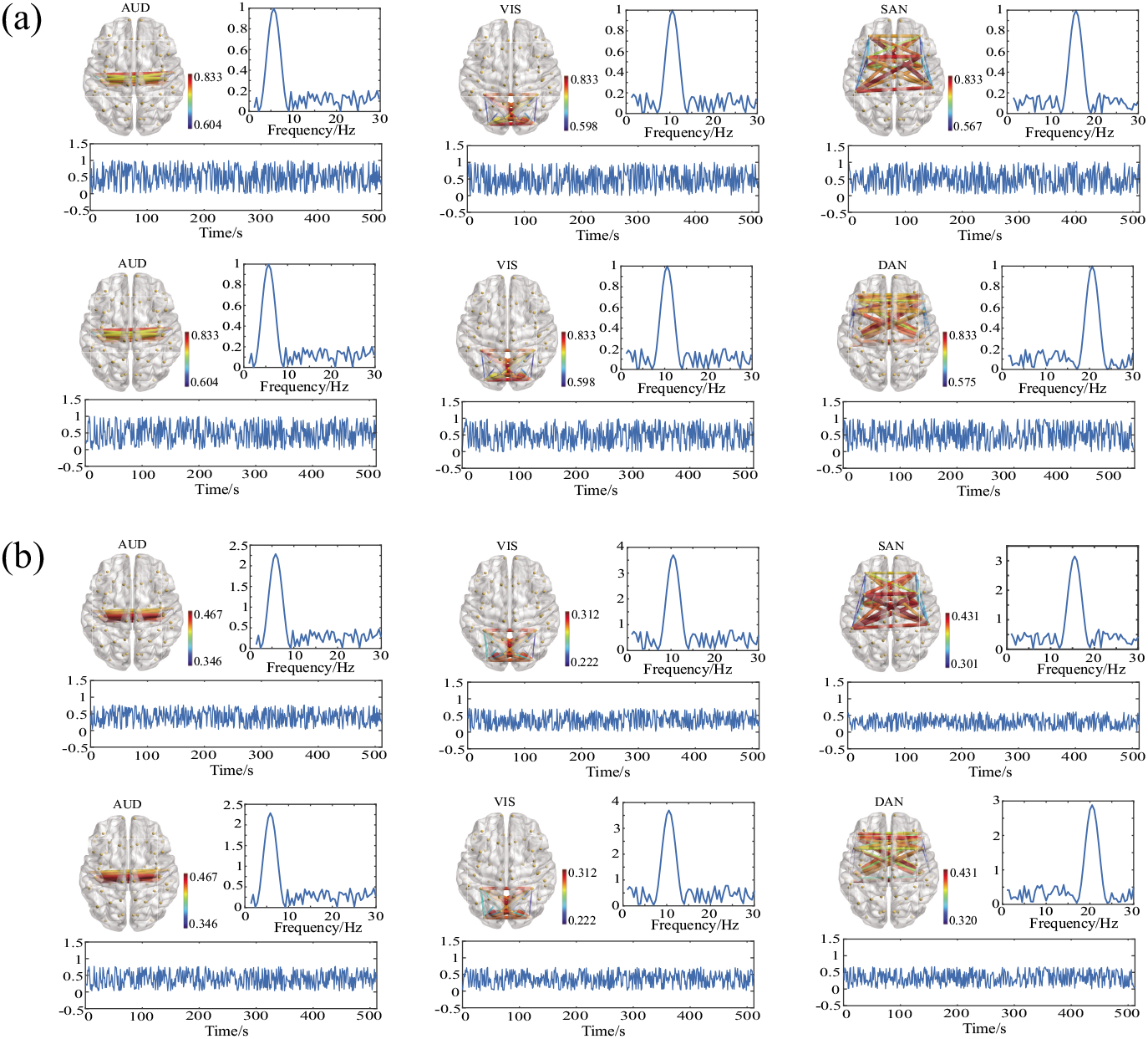
Simulation illustration. (a) Three spatio-temporal-spectral patterns were simulated for two groups, and the first two patterns were identical in adjacency and spectral modes. (b) The reconstructed spatio-temporal-spectral patterns.

#### 2) simulation results

We implemented the low-rank DC-NTD-FHALS algorithm on the synthetic data, and we set *SNR* = 15, *L*_*f*_ = *L*_*c*_ = 2, and *R*_HC_ = *R*_MDD_ = 3. The extracted temporal, spectral and adjacency factors were shown in Figure 2(b). We ran 10 times of the low-rank DC-NTD-FHALS algorithm, and we obtained stable decomposition results with an averaged tensor fit of 0.864 and an averaged running time of 113.27 seconds.

### B. Results of EEG data

Through PCA analysis on the unfolded data along the spectral mode for two-block data, we extracted *R*_*HC*_ = *R*_*MDD*_ = 27 components for both HC group and MDD group. Then we performed the fourth-order CP tensor decomposition on each block data, and computed the correlation maps of spectral and adjacency factors between two groups, as shown in Figure 3. According to the correlation maps, we set the number of coupled components in the spectral mode *L*_*f*_ = 25 and the number of coupled components in the adjacency mode *L*_*c*_ = 7. We ran 20 times of the low-rank DC-NTF-FHALS algorithm, and the averaged running time was 12616 seconds. The running time was 63819 seconds by one implementation of the DC-NTF-FHALS algorithm without the low-rank approximation, which indicated that the low-rank approximation could greatly reduce the computational load.

**Fig. 3:**
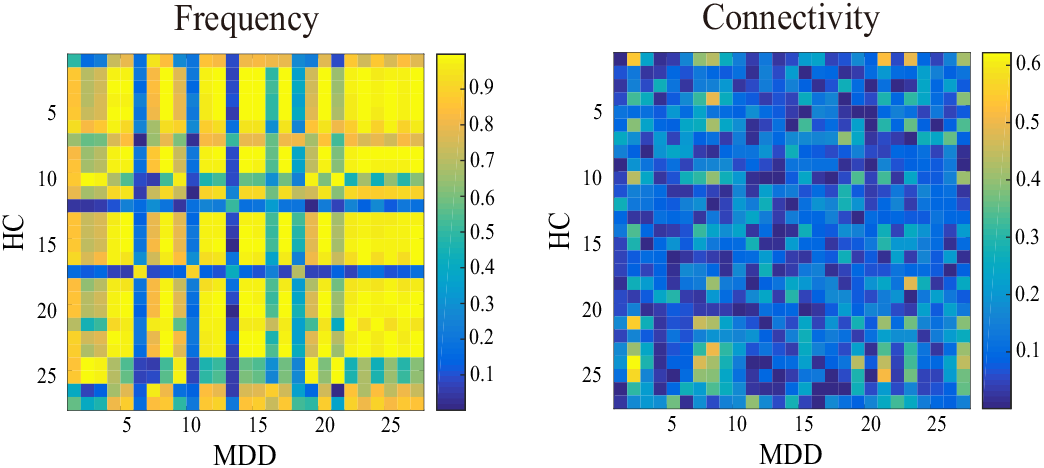
Correlation analysis of the spectral factor and the adjacency factor extracted from the 4-D tensor decomposition for each block data.

After applying the low-rank DC-NTF-FHALS algorithm and correlation analysis with musical features, we summarized the results of 20 times of implementation and obtained three oscillatory hyperconnectivity networks, as shown in Figure 4, and three oscillatory hypoconnectivity networks, as shown in Figure 5. For hyperconnectivity networks, Figure 4I shows a right hemisphere dominated network modulated by oscillations of alpha and beta (10-16 Hz) bands and the musical feature of Fluctuation Centroid. The strong connections of this network connect the core regions of DMN, including medial prefrontal cortex (mPFC), precuneus cortex, and posterior cingulate cortex (PCC). Figure 4II indicates a left auditory-related network modulated by delta oscillations and the Fluctuation Centroid feature. An aberrant delta-specific prefrontal network is identified, which is related to the musical feature of Key Clarity, as shown in Figure 4III. For hypoconnectivity networks, Figure 5I and Figure 5II exhibit fronto-parietal networks which are mainly related to attention control. The fronto-parietal networks are modulated by oscillations of 8-14 Hz and 10-19 Hz and musical features of Mode and Fluctuation Entropy, respectively. Figure 5III shows a low-frequency (delta oscillations) modulated prefrontal network which is significantly related to the musical feature of Mode, and this network is implicated in complex cognitive functions.

**Fig. 4:**
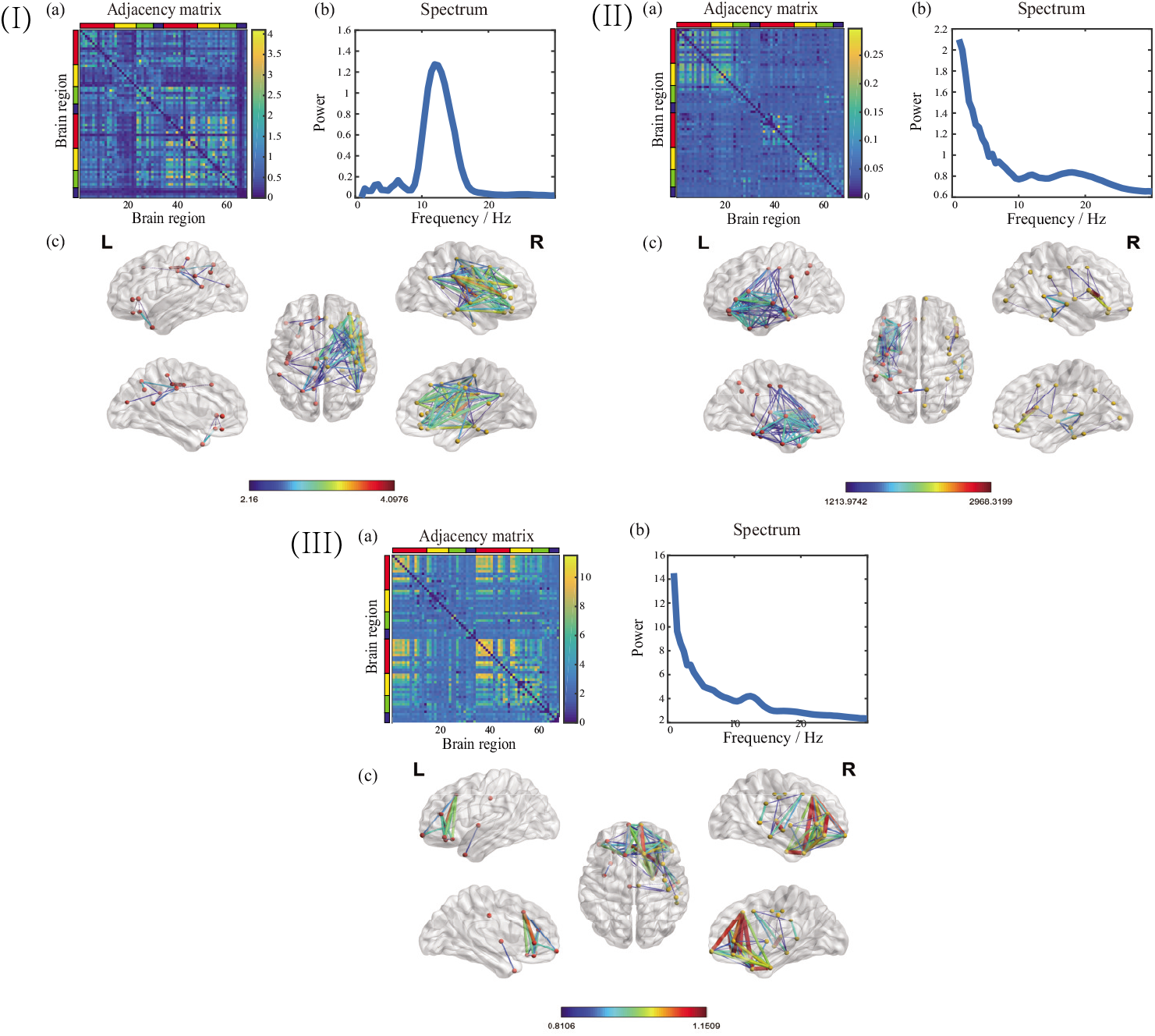
Three oscillatory hyperconnectivity networks. (a) Adjacency matrix representation of the network. The 68 brain regions are ordered from the left hemisphere to the right hemisphere. Within each hemisphere, the brain regions are arranged in the order of frontal lobe, temporal lobe, parietal lobe, and occipital lobe, as indicated in red, yellow, green, and blue color, respectively. Within each lobe, the brain regions are ordered according to their y-location from anterior regions to posterior regions. (b) The spectral component of the network. (c) Cortical space representation of the network in Lateral, medial and dorsal view. The networks I, II, and III are related to the musical features of Fluctuation Centroid, Fluctuation Centroid, and Key Clarity, respectively.

**Fig. 5:**
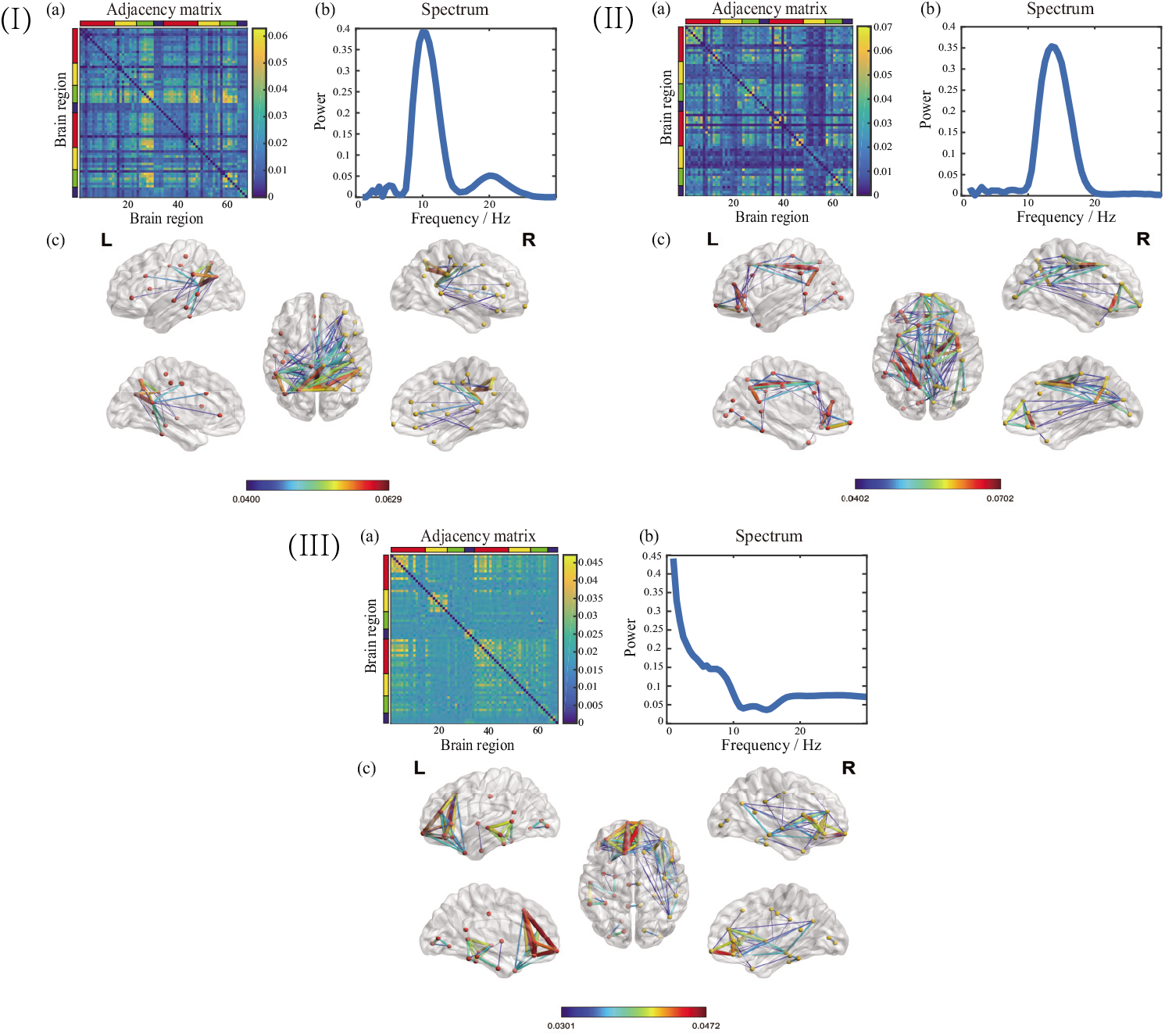
Three oscillatory hypoconnectivity networks. (a) Adjacency matrix representation of the network. The 68 brain regions are ordered from left hemisphere to right hemisphere. Within each hemisphere, the brain regions are arranged in the order of frontal lobe, temporal lobe, parietal lobe, and occipital lobe, as indicated in red, yellow, green, and blue color, respectively. Within each lobe, the brain regions are ordered according to their y-location from anterior regions to posterior regions. (b) The spectral component of the network. (c) Cortical space representation of the network in Lateral, medial and dorsal view. The networks I, II, and III are related to the musical features of Mode, Fluctuation Entropy, and Fluctuation Entropy, respectively.

## IV. Discussion

As far as we know, this study is the first attempt to investigate the aberrant dFC across temporal evolution and spectral modulation in MDD during music listening based on a coupled tensor decomposition approach. Fully considering the high-dimensional data structure and group differences, we proposed a comprehensive framework to extract the FC networks characterized by spatio-temporal-spectral modes of covariation. We identified the common oscillatory networks shared among HC and MDD groups and the individual oscillatory networks specified in the MDD group. Then, we summarized three overactive functional networks and three underactive functional networks according to the analysis of musical modulations. We also verified our method using simulated data.

MDD is characterized with imbalanced communications among large-scale functional networks, including hyperconnectivity and hypoconnectivity within specific brain networks or between distinct brain networks during resting-state, see a meta-analysis in study [6]. In our study, we also found hyperconnectivity and hypoconnectivity functional networks in naturalistic music perception. We identified a right hemisphere dominated hyperconnectivity network which involved the essential regions of DMN, including mPFC, PCC and precuneus cortex, as shown in Figure 4I. The hyperconnectivity in DMN are often considered as reflecting rumination, where MDD patients perseverate on negative, self-referential thoughts [49], [50]. Many researches have reported the hyperconnectivity within DMN in MDD, which supports that within-DMN hyperconnectivity is related to enhanced the positive connectivity in MDD [6], [50]. Figure 4II shows a delta band-modulated and left auditory-related network, which is activated by a rhythmic feature of Fluctuation Centroid. The delta band was demonstrated to have a substantial influence on the identification of natural speech fragments in a MEG study [51], and the decoding of rhythmic features was found to be significantly correlated with the auditory cortex during music perception [20], [52]. The abnormal delta band-modulated and left auditory-related network identified in our study might indicate that MDD patients were less involved in music perception. We identified two delta band modulated prefrontal networks, both of which were related to tonal features, Key Clarity and Mode, as shown in Figure 4III and Figure 5III. However, the prefrontal network in Figure 4III was hyperactive and right hemisphere lateralized, and the prefrontal network in Figure 5III was hypoactive and left hemisphere lateralized. The prefrontal cortex has been implicated in planning complex cognitive behavior, decision making and working memory. There are numerous lines of evidence demonstrating that prefrontal cortex is dysregulated in depression, and both increased and decreased functional connections in the prefrontal network may lead to the failure of inhibitory control in depression [53]–[56]. Those two prefrontal networks also have abnormal connections with temporal poles, which may indicate the dysfunction in semantic integration during music listening [57], [58]. Our findings are well supported by those literatures that the dysconnectivity in the prefrontal network can influence the high-order cognitive functions and information integration during music perception in MDD. Figure 5I and Figure 5II indicate hypoconnectivity fronto-parietal networks modulated by different oscillations and musical features. The abnormal development of the fronto-parietal network is a common feature across many psychiatric disorders with the deficit in cognitive control. Previous studies have demonstrated that MDD is characterized by hypoconnectivity within the frontoparietal network, which is involved in the top-down modulation of attention and emotion [6], [18], [59].

The proposed approach in this study is a totally data-driven technique, which detects the oscillatory brain networks spanning to the whole brain and avoids the selection of regions of interest according to prior knowledge. In our study, the key issue of applying coupled tensor decomposition is the selection of the number of all the extracted components and the number of coupled components. There are several methods for the selection of the number of extracted components in tensor/matrix decomposition, such as PCA, the difference of fit (DIFFIT), model order selection, and so on [30]. In our study, due to that the spectral mode retains the minimum samples compared with temporal and adjacency modes, we used PCA applied on the unfolded data along the spectral mode to determine the number of extracted components. We believe this unfolding format can help to approach the true underlying low-dimensional space. However, the selection of the number of coupled components and the coupled modes mainly relys on the data characteristics used in the applications. Refer to our previous study, we use a correlation analysis in the spectral and adjacency modes in our study [24].

The scales of the reconstructed spatial, temporal and spectral factors are different from those in the synthetic data, see Figure 2. This is because that we did not add constraints on the scales of factors in the implementation of the low-rank DC-NTF-FHALS algorithm, but the scales are the same after back-projection of the factors. The scale indeterminacy will not change the topology of networks, the evolution of time courses and the modulation of oscillations. However, the addition of the constraints on scales will increase the model complexity and computational cost. In the present study, we only consider the group differences between HC and MDD groups by extracting the common features shared by two groups and individual features owned by each group. Subject differences are omitted, and we assume that the extracted components are shared by all the subjects within each group. The subject differences are covered in the residuals of the coupled tensor decomposition, which are not concerned in this research. The problem of subject differences may bring more challenges, like statistical analysis, but it is also a crucial and realistic issue, especially in clinical applications.

## V. Conclusion

In this study, we investigated the oscillatory hyperconnectivity and hypoconnectivity networks elicited by musical stimuli in MDD. Considering the high-dimensional structure of the datasets and group differences between HC and MDD groups, a comprehensive framework was proposed based on coupled tensor decomposition, and six abnormal connectivity networks with spatio-temporal-spectral modes of covariation were identified in MDD during music listening. Our findings are well supported and verified by previous literatures. Our research may serve as a signature of the brain’s functional topography characterizing MDD, and provide novel biomarkers for the clinical diagnosis and treatment in MDD. The spectral profiles and spatial networks are usually characterized with sparsity, and the sparse regularization will be considered in the coupled tensor decomposition model in the future work. The neural correlates and dynamic neural processing of musical emotions have not been well studied, and the future work will also focus on the selection of control stimuli.

